# Detection and Annotation of Unique Regions in Mammalian Genomes

**DOI:** 10.1101/2024.10.11.617789

**Authors:** Beatriz Vieira Mourato, Bernhard Haubold

## Abstract

Long unique genomic regions have been reported to be highly enriched for developmental genes in mice and humans. In this paper we identify unique genomic regions using a highly efficient method based on fast string matching. We quantify the resource consumption and accuracy of this method before applying it to the genomes of 18 mammals. We annotate their unique regions of at least 10 kb and find that they are strongly enriched for developmental genes across the board. When investigating the subset of unique regions that lack annotations, we found in the tasmanian devil the gene encoding iniositol polyphosphate-5-phosphatase A, which is an essential part of intracellular signaling. This implies that unique regions might be given priority when annotating mammalian genomes. Our documented pipeline for annotating unique regions in any mammalian genome is available from the repository github.com/evolbioinf/auger; additional data for this study is available from the data-verse at doi.org/10.17617/3.4IKQAG.

## Introduction

Transposons were discovered in the mid 1940’s by Barbara McClintock, a few years before the discovery of the double-helical structure of DNA by Watson and Crick in 1953. The double helix focused attention on the linear sequence of its nucleotides, while transposons suggest this linearity can be altered. McClintock observed that the moblilization of transposons often follows unexpected shocks to a cell, for example through double strand breaks, infection by retroviruses, or hybridization (McClintock, 1984). McClintock further noted that transposon activation in somatic cells can lead to changes in gene regulation.

When the human genome was first published, its transposon content, roughly 50 %, could be quantified with unprecedented precision (Intern. HG Seq. Cons., 2001). It was noted at the time that the *Hox* clusters had the lowest transposon density of any region of similar size in the genome. The *Hox* clusters are approximately 100 kb long and contain about ten transcriptional regulators that determine an organism’s basic body plan. Mammals have four *Hox* clusters, *HoxA, HoxB, HoxC*, and *HoxD*. The authors of the human genome paper speculated that selection against changes in the regulation of *Hox* genes kept these regions free of transposons.

This observation sparked further interest in the functional content of a newly defined category of DNA sequence, transposon-free regions (TFRs) (Simons et al., 2005). Apart from containing no transposons, TFRs also contained no satellite DNA, and no more than 20 % of their length was homologous to other parts of the genome. This left approximately 1000 regions of at least 10 kb in the genomes of human and mouse covering a dozen Mb, of which 90 % were non-coding. The largest TFR in the human genome was 81 kb long and intersected *HoxA*. In general, TFRs were highly enriched for developmental genes and transcription factors (Simons et al., 2005).

Of the two possible mechanisms for maintaining TFRs, exclusion and purifying selection, selection is generally favored (Intern. HG Seq. Cons., 2001; Simons et al., 2005). This suggests that mammalian genomes can be read as the result of transposon mutagenesis experiments run over evolutionary time.

A year after the first description of TFRs, a novel chromatin state was discovered, characterized by the co-occurrence of activating H3K4me3 marks and repressing H3K27me3 marks (Bernstein et al., 2006). These “bivalent regions” are found in the chromatin of embryonic stem cells where they are associated with developmental genes; they vanish as cells differentiate. Moreover, bivalent regions are located in TFRs.

Bivalency has since attracted a lot of attention as it is associated not only with cell differentiation, but also with its opposite, cell de-differentiation, better known as cancer. The distribution of transposons, on the other hand, is often less well studied (Platt II et al., 2016). Moreover, TFRs are not simply the complement of transposons, but so far require for their identification a combination of repeat detection and searches for self-homology.

The best-known tool for repeat detection is Repeat-Masker, which takes as input a genome sequence and a set of transposon sequences; it then marks the location of the transposons in the genome. On the scale of mammalian genomes this amounts to a considerable number of homology searches and hence computational effort. However, the toolset of genomics has changed since the inception of RepeatMasker almost thirty years ago (Smit, 1996) . For example, fast exact matching using suffix trees or suffix arrays has become more accessible through new algorithms (Abouelhoda et al., 2002) and libraries (Fischer and Kurpicz, 2017). This has resulted in read mappers like Bwa (Li and Durbin, 2009) and genome aligners like Mummer (Marçais et al., 2018).

Fast suffix array construction also underlies the program Macle for quickly detecting TFRs (Pirogov et al., 2019). Macle, which stands for *match complexity*, is based on maximal matches, that is, matches that cannot be extended without losing the match. Let *m*_o_ be the observed number of maximal matches that intersect a sliding window on a genome, and *m*_e_ its expectation for random sequences, then the *match complexity* is defined as

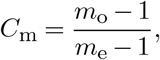

where the subtraction of 1 ensures that *C*_m_ ranges from zero to an expectation of one. Given the null distribution of *C*_m_ (Pirogov et al., 2019), regions where *C*_m_ is indistinguishable from random can be picked. These regions have no homologs elsewhere in the genome and hence are free of transposons. However, the regions detected with Macle are also free of any other types of repeats, while they may conceivably contain a single-copy transposon. Hence we call them *unique regions*, URs.

With a sliding window of 10 kb, the human genome contains 1243 URs covering 17.3 Mb and 772 URs in the mouse genome covering 10.1 Mb (Pirogov et al., 2019). These are similar numbers to those found earlier for TFRs in human and mouse. Moreover, enrichment of URs for developmental genes was greater than 10-fold and highly significant in both organisms (Pirogov et al., 2019).

Here we extend the analysis of URs to a sample of 18 mammalian genomes from nine orders covering placentals and marsupials. We identify the URs in these genomes and test for functional enrichment. This requires maps between gene identifiers and GO terms for organisms other than human and mouse, where they are readily available. We present a computational workflow for constructing such maps from scratch for any sequenced mammal. Given these maps, they are used in a Monte Carlo test for enrichment. In agreement with previous studies, we find in all 18 genomes studied that URs are highly enriched for developmental genes. This establishes a strong link in mammals between a now easily observable property of DNA sequences— uniqueness—and function. In addition, we investigate URs that do not intersect any annotations, *anonymous* URs. The longest anonymous UR turns out to intersect the gene encoding a well-known component of intracellular signaling.

## Material & Methods

### A. Software

All software repositories mentioned in the following sections are located at

github.com/evolbioinf

### B. Comparing Macle and RepeatMasker

We compared Macle version 0.1 and RepeatMasker version 4.1.5. Macle runs on a single thread. Repeat-Masker was run with 10 threads. Both programs were applied to sequence data containing mutations simulated with the program Mutator, which is part of the Biobox repository. Macle was run with 1 kb windows and RepeatMasker with default parameters. Macle runs on an index of the input file. The 18 Macle indexes used throughout this study are available from the accompanying dataverse.

### C. Sequence Data and Phylogeny

We picked mammalian orders for which we found at least two genomes assembled to chromosome level. This resulted in a sample of nine orders and hence 18 genomes, which are listed in Supplementary Table S1. We used the program Datasets supplied by the NCBI to download these genomes, their GFF3 annotation files, and their proteomes. We describe the details of how to download the data and to extract unique regions in our repository Auger, for “analyze unique genomic regions”. We calculated the phylogeny of the 18 genomes sampled from their *HoxD* regions, which were aligned with Mafft (Katoh et al., 2005). The alignment was subjected to maximum likelihood phylogeny reconstruction with Iqtree2, which we used to search for the best-fitting model and to bootstrap it 1000 times in slow mode (Kalyaanamoorthy et al., 2017).

### D. Functional Enrichment

Functional enrichment was assessed using the programs Annotate, Ego, and Shuffle from our Gin repository. The program Annotate takes genomic intervals extracted with Merwin (Auger repo) from Macle output and a GFF3 file as input and returns a list of genes whose promoters intersect the intervals. Promoters were defined as 2 kb intervals upstream of transcription start sites (Robertson et al., 2007). Instead of promoters, Annotate can also intersect transcripts. The program Ego was used to count the number of genes per GO term.

The graph of enriched GO terms was drawn using the “chart” link of the QuickGO REST API at www.ebi.ac.uk/QuickGO/api/index.html

#### D.1. Monte Carlo Test

We used a Monte Carlo test to assess the significance of the gene counts per GO term. Only GO terms with at least ten genes were included in the analysis. The program Shuffle (Gin repo) was used to shuffle the unique intervals across the genome of origin. Then we annotated them again and counted the frequency with which a gene count at least equal to the original count was observed. This frequency is the desired *P* -value, the error probability when rejecting the null hypothesis that the observed gene count is due to chance. The test was carried out with one million iterations and the resulting *P* -values were Bonferroni-corrected by multiplication with the number of tests, that is, the number of GO terms with at least ten genes. GO terms with corrected *P* ≤ 0.01 were considered enriched.

#### D.2. Mapping Gene IDs to GO Terms

The enrichment analysis just described depends on a map between the gene IDs extracted from the GFF3 file and the GO terms that give them functional meaning. This map is provided by the file `gene2go` supplied by the NCBI:

ftp.ncbi.nih.gov/gene/DATA/gene2go.gz

The `gene2go` file originally consists of eight columns, of which the four columns shown in Figure 1A are used in our analysis. These are Gene ID (red), GO ID (green), GO term (blue), and Category (also blue). Unfortunately, the original `gene2go` file contains only GO terms for six of our 18 mammalian species. So we constructed our own `gene2go` files. To make results comparable across species, we carried out this construction even for the six species represented in the original `gene2go` file.

**Figure 1.**
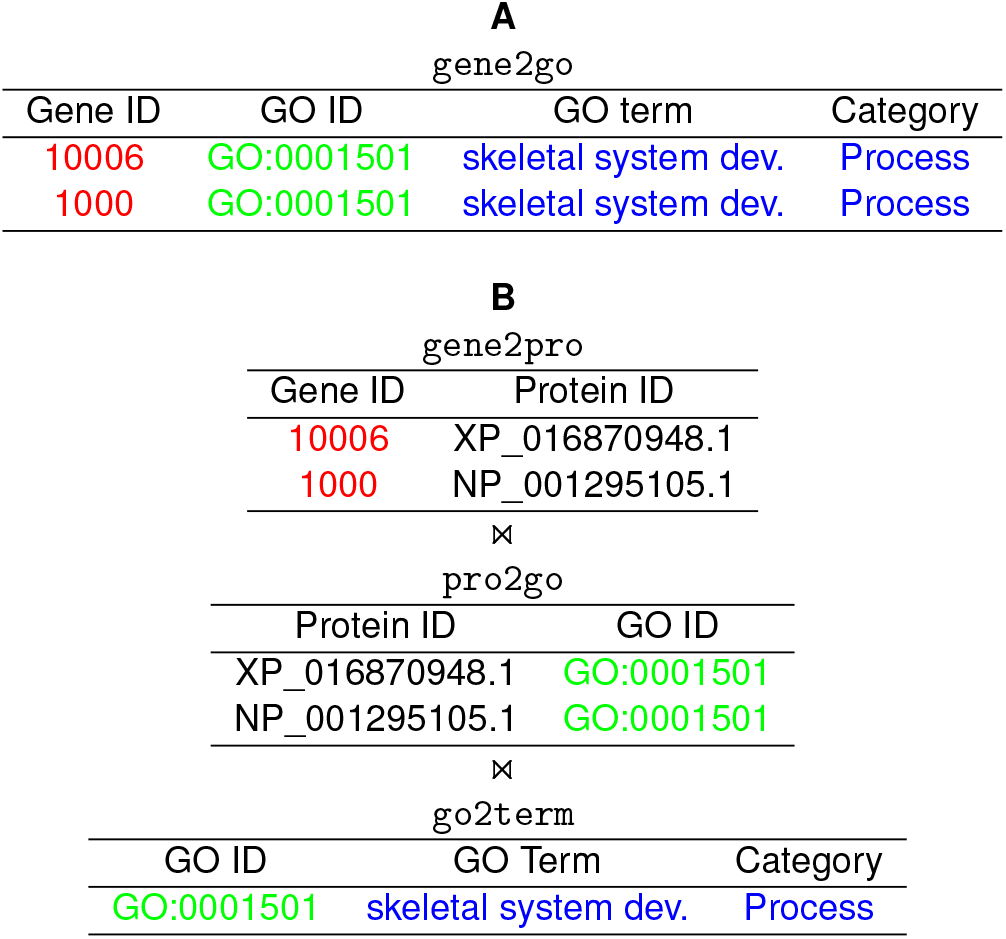
Sample `gene2go` file (A), and its construction (B) by joining (⋉⋊) the three tables `gene2pro, pro2go`, and `go2term`; colors indicate corresponding columns.

As shown in Figure 1B, the `gene2go` file is constructed by joining three tables. The table gene2pro maps gene IDs to protein IDs and is extracted from the GFF3 file. The table `pro2go` maps protein IDs to GO IDs and is constructed by subjecting the proteome to homology analysis with the `eggNOG` software (Cantalapiedra et al., 2021). The table `go2term` maps GO IDs to GO terms and categories, and is extracted from the current GO database distributed by the GO consortium via their website geneontology.org; it can be downloaded from

https://purl.obolibrary.orgøbo/go/go-basic.obo

Like the data handling, the enrichment analysis and the construction of the `gene2go` files is described in the Auger repo. The 18 `gene2go` files used in this study are also available from the accompanying dataverse.

## Results

### A. Comparing Macle and RepeatMasker

We begin by comparing our tool for picking unique regions, Macle, to an established tool for marking repeats, RepeatMasker. For the purpose of this comparison we treat picking unique regions as complementary to picking repeats. We focus on the sensitivity of the respective tools, which we quantify through simulation and application to the human genome.

For the simulation, we analyze two identical copies of the human transposon MER5A1r, which is 3.7 kb long, on the background of a random 25 kb sequence. Figure 2 shows the proportion of the transposon identified by Macle with 1 kb sliding windows and Repeat-Masker as a function of the mutation rate. For Macle that proportion declines sharply at 0.2 mutations/site, while for RepeatMasker this happens much later at 0.5 mutations/site.

**Figure 2.**
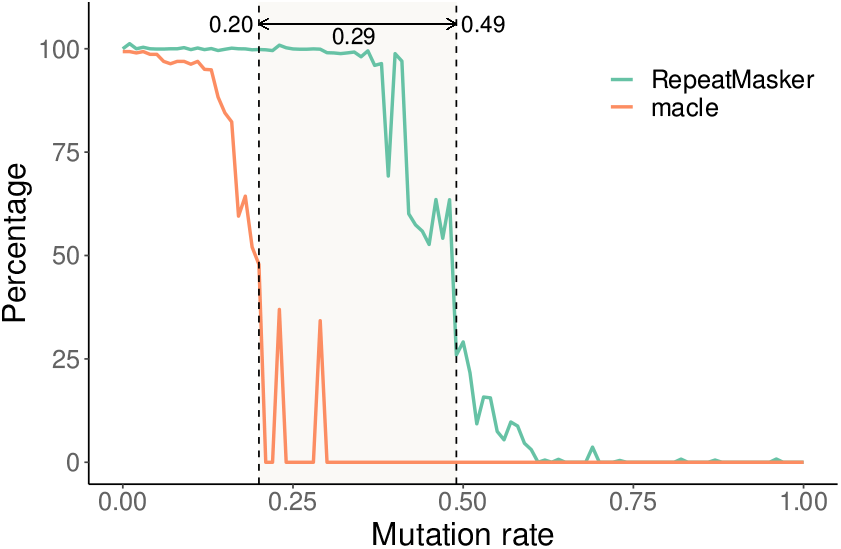
Detecting two identical repeats as a function of mutation rate. Percentage of nucleotides masked by RepeatMasker, and percentage of non-complex nucleotides calculated by Macle, for a given mutation rate. Vertical lines denote the mutation rate where the percentage first fell below 50 %.

To compare these simulation results to real data, we applied Macle again with 1 kb windows and RepeatMasker to the human genome. For a start, this revealed the quite different resource requirements of the two programs. Macle took 1 h 14 m and 218.0 GB RAM to index the human genome; parsing this index took another 36.0 s and 26.9 GB of memory. In contrast, Repeat-Masker required 44 h 19 m and 80.1 GB RAM to mask the human genome, that is, over 30 times longer than the indexing step of Macle but occupying less than half the memory.

Having completed the Macle and RepeatMasker runs on the human genome, we looked up the intersection between masked and unique regions. From our simulations in Figure 2 we expected this overlap to be made up of sequences whose closest homolog in the genome has diverged by more than 0.2. Figure 3 shows the divergence of repeats missed by Macle as a function of their divergence rate for five categories of repeat lengths. As expected, the divergence of these regions tends to be greater than 0.2 mutations/site, unless they are shorter than 100 bp. The exception to this rule is the single region in the bottom row of Figure 3. It originates from a single copy of the 7.2 kb human endogenous virus HERV-Fc2 that has diverged by 16.9 % from the consensus sequence.

**Figure 3.**
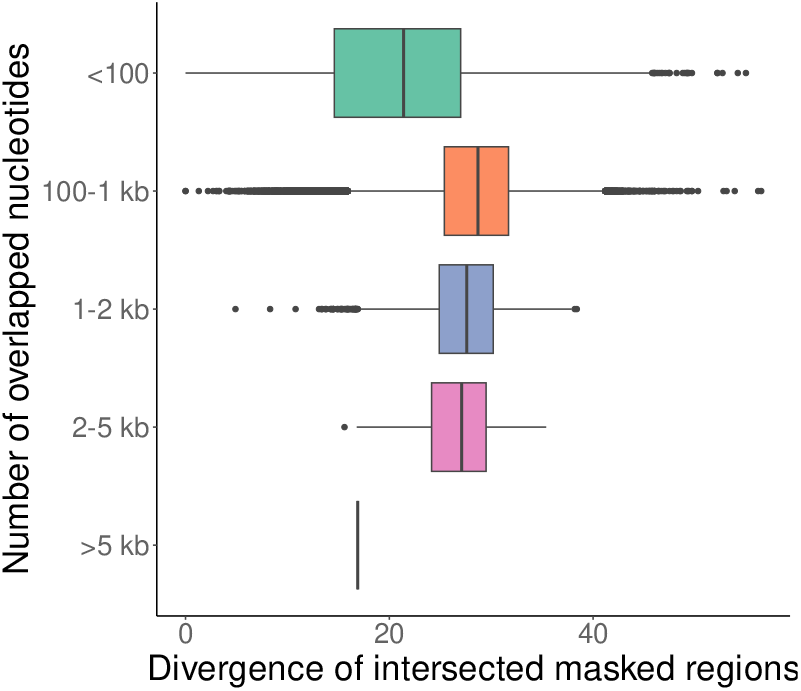
Comparing the divergence rates for overlaps between unique regions and repeats for overlaps ranging from less than 100 bp to greater than 5 kb.

### B. Unique Regions in Mammalian Genomes

After establishing that the unique regions identified by Macle are largely free of recent transposon insertions, we proceeded to download the genomes of our 18 target organisms. Figure 4 shows their phylogeny calculated from the *HoxD* cluster. Note the deep split between marsupials and placentals, and the two super-orders of the Laurasiatheria and the Euarchontoglires.

**Figure 4.**
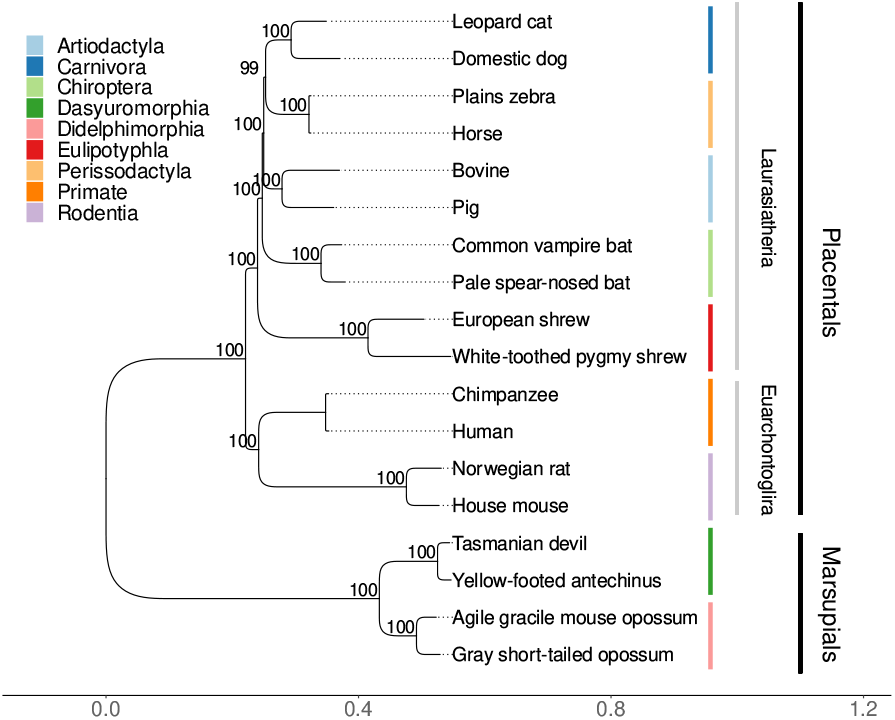
Bootstrapped maximum likelihood tree of the *HoxD* cluster of the 18 species analyzed, 12 placentals and 4 marsupials.

In order to pick unique regions from the downloaded genomes with Macle, they need to be indexed first. Figure 5A shows that the run time of indexing is roughly linear in the genome size, which ranged from 2.1 Gb for the bats (Chiroptera) to 3.7 Gb for the agile gracile mouse opossum (Didelphimorphia), leading to user times between 1983.3 s and 4056.0 s. Similarly, the memory consumption of indexing is linear in the genome length and ranges from 139.8 GB to 238.1 GB (Figure 5B).

**Figure 5.**
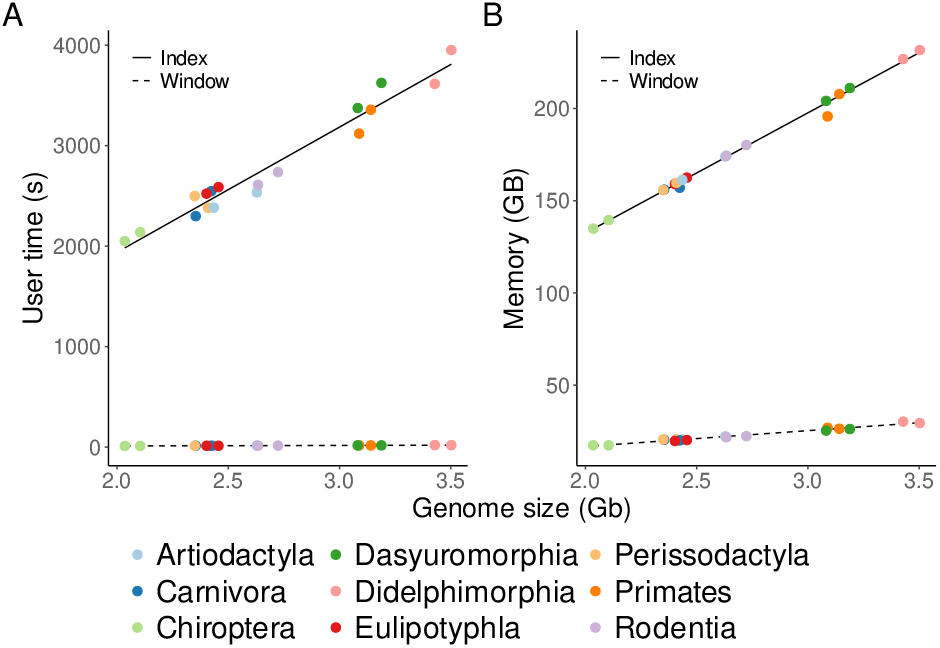
Time (A) and memory (B) consumption of Macle as a function of genome size when calculating an index (*Index*) or when carrying out a sliding window analysis on an index (*Window*).

Given these indexes, their analysis is much less resource intensive with user times ranging from just 11.8 s to 19.8 s and memory requirements from 17.4 GB to 30.3 GB (Figure 5). Since using a Macle index is over 170 times faster than computing it, we make the 18 indexes constructed for this study available as part of the accompanying dataverse.

### C. The Effect of Window Size

To explore the effect of window size on our analysis, we varied the window size from 1 kb to 50 kb. Figure 6A shows that the proportion of unique regions (URs) declines sharply as a function of window length. For 1 kb windows the fraction of URs ranges from 34.4 % for the leopard cat (Carnivora) to 10.8 % for the european shrew (Eulipotyphla). Compare this to the 49.1 % of the human genome masked by RepeatMasker, which might imply that the complement, 50.9 %, is unique. However, according to Macle with 1 kb sliding window, only 13.5 % of the human genome is unique, because Macle recognizes any repeat, not just the repetitive elements collected in a database. As the window size increases, the fraction of URs declines to between zero in the european shrew and 0.6 % in the tasmanian devil.

**Figure 6.**
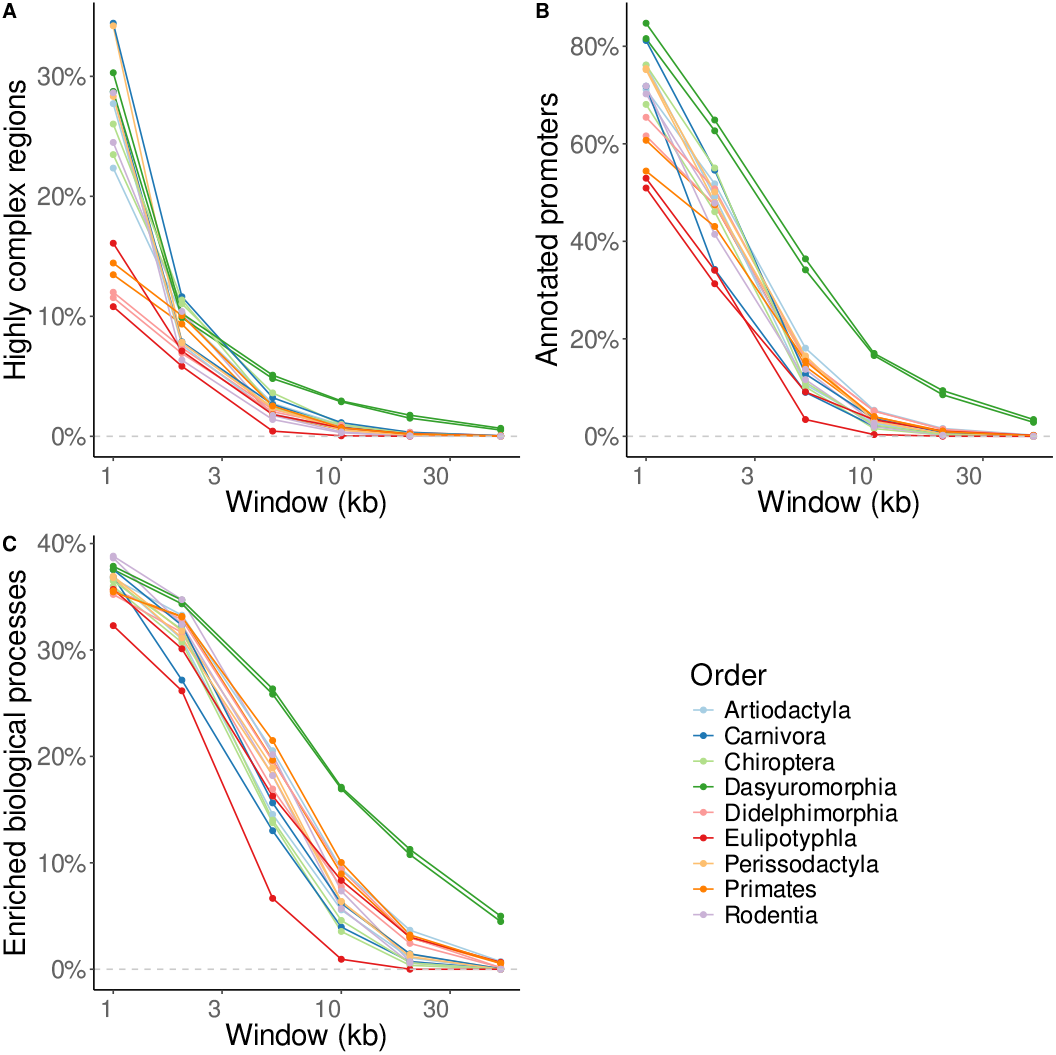
Highly complex regions in 18 mammalian genomes as a function of sliding window size. A) Total length; B) Annotated promoters; C) Enriched biological processes.

Similarly, the fraction of promoters that intersect URs declines with increasing window length. For 1 kb windows, the fraction of intersecting promoters ranges from 84.7 % for the tasmanian devil (Dasyuromorphia) to 50.9 % for the white-toothed pigmy shrew (Eulipotyphla) (Figure 6B). With 10 kb windows this is down to an average of 4.7 %, with a range from 0.4 % for the european shrew to 17.0 % for the tasmanian devil.

Likewise, the number of enriched GO terms decreases continuously with increasing window length. As shown in Figure 6C, for 1 kb windows the fraction of enriched GO terms varies between 32.3 % for the european shrew (Eulipotyphla) and 38.8 % for house mouse (Rodentia). For 50 kb windows this declines to values between zero in eight species (pig, domestic dog, bats, european shrew, plains zebra and rodents) and 5.0 % in the tasmanian devil (Dasyuromorphia).

Doubling the window size to 2 kb slightly decreases the number of enriched GO terms; now 26.2 % of terms are enriched in the european shrew, and 34.7.1 % in the house mouse.

Having surveyed the fraction of enriched GO terms as a function of window length, we now turn to their enrichment ratios. Figure 7 shows the average enrichment ratios as a function of window length for each of the nine orders. The curves for the two species analyzed per order are quite similar. However, the enrichment ratios do not behave uniformly across orders. In almost all of the orders the average enrichment ratio plateaus as a function of window length. In contrast, in the Eulipotyphla and Primates the average enrichment ratio grows continuously with window size.

**Figure 7.**
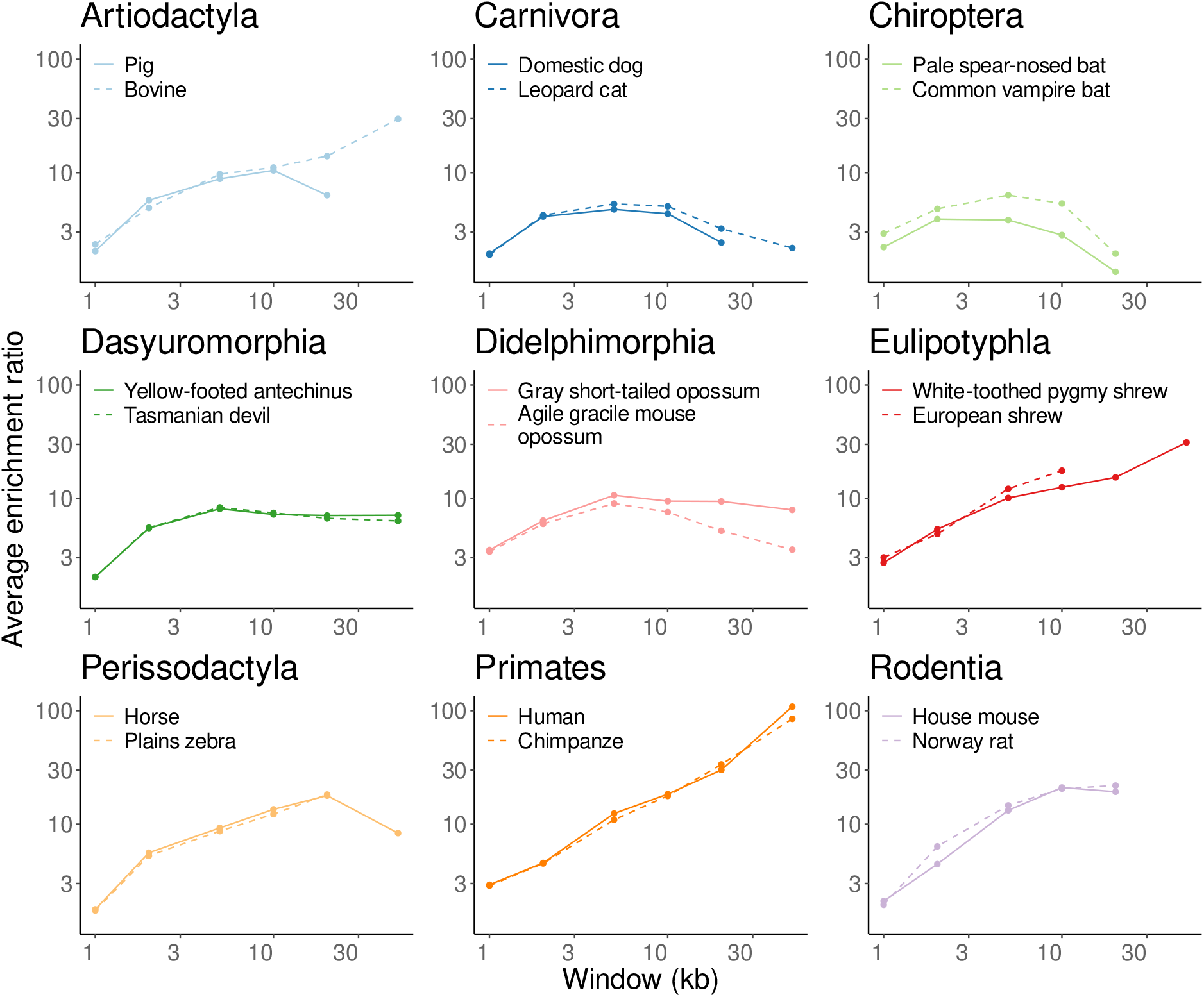
Average enrichment ratio per genome as a function of window size for the nine mammalian genera investigated.

### D. Enrichment of GO Terms

The next question we address is, which GO terms are enriched? For this we restrict our analysis to 10 kb windows, though qualitatively it makes little difference whether we choose 5 kb, 10 kb, or 20 kb as window length (not shown). We concentrate on the GO terms enriched in all 18 species studied.

Table 1 shows the ten most enriched of these terms, all of which explicitly refer to development, ranging from “Anterior/posterior pattern specification” with an enrichment ratio ranging from 6.8 in the pale spear-nosed bat to 89.1 in the european shrew, to “Chordate embryonic development” with an enrichment ratio ranging from 4.2 in the pale spear-nosed bat and 31.5 in the norwegian rat.

**Table 1.** Enriched biological processes shared across the 18 mammalian species investigated. Top 10 processes with the highest overall enrichment ratio are shown. The respective enrichment ratio for each species for each term is shown. The darker the hue, the higher the enrichment ratio.

|  | Artiodactyla |  | Carnivora |  | Chiroptera |  | Dasyuromorphia |  | Didelphimorphia |  | Eulipotyphla |  | Perissodactyla |  | Primates |  | Rodentia |  |
| --- | --- | --- | --- | --- | --- | --- | --- | --- | --- | --- | --- | --- | --- | --- | --- | --- | --- | --- |
| Anterior/posterior pattern specification | 32.1 | 27.6 | 12 | 12.5 | 8.9 | 6.8 | 13.1 | 12.5 | 22.3 | 18.3 | 89.1 | 33.7 | 40.9 | 36.2 | 50.6 | 45.8 | 60.5 | 54.2 |
| Regionalization | 31.7 | 27 | 10.4 | 12.6 | 11.5 | 6.3 | 12.2 | 12.3 | 23.1 | 17.2 | 62 | 33.7 | 41.4 | 36.9 | 50.8 | 44.7 | 55.9 | 53.2 |
| Pattern specification process | 27.7 | 23.6 | 10.3 | 11.5 | 10.9 | 5.8 | 11.1 | 11.3 | 21.4 | 16.1 | 50.4 | 27.7 | 36.7 | 32.3 | 44.4 | 39 | 48.2 | 44.5 |
| Embryonic organ development | 23.1 | 20.4 | 8.8 | 9.5 | 10.5 | 6 | 10.3 | 10.7 | 18.7 | 13.5 | 65.9 | 24.4 | 30.2 | 27.1 | 36.5 | 31.5 | 38.2 | 35 |
| Embryonic morphogenesis | 22.4 | 21.1 | 8.8 | 9.3 | 10.2 | 5.9 | 9.8 | 10.3 | 17.8 | 13.9 | 51.4 | 22.8 | 31.1 | 26.6 | 37.9 | 31.9 | 39.6 | 38.8 |
| Forebrain development | 19.8 | 18.1 | 6.6 | 7.9 | 10.9 | 5.2 | 9.2 | 9.8 | 14.7 | 9.6 | 61.3 | 21.9 | 25.9 | 25.1 | 35.7 | 34 | 35.6 | 32.5 |
| Animal organ morphogenesis | 20 | 16 | 7.7 | 7.9 | 8.6 | 4.8 | 8.9 | 9 | 16 | 11.4 | 29.1 | 19.6 | 25.1 | 22.4 | 31.9 | 27.7 | 31.1 | 28.1 |
| Negative regulation of transcription by RNA polymerase II | 20.7 | 14.8 | 6 | 6 | 7.2 | 3.8 | 9.6 | 10.1 | 14.5 | 11 | 44.9 | 17.8 | 25.4 | 21.1 | 26.7 | 25.3 | 30.7 | 27.9 |
| Brain development | 18.4 | 15.2 | 5.9 | 7.2 | 8 | 4.1 | 9 | 9.7 | 13.9 | 10.3 | 35.6 | 19.7 | 21.9 | 19.9 | 29.4 | 27.5 | 32.3 | 31.5 |
| Chordate embryonic development | 19.8 | 15.9 | 7 | 7.2 | 7.6 | 4.2 | 9.2 | 9.7 | 15.2 | 12.5 | 26.8 | 19.9 | 24.9 | 20.8 | 29.6 | 24.8 | 30.9 | 31.5 |
|  | Bovine | Pig | Domestic dog | Leopard cat | Common vampire bat | Pale spear-nosed bat | Yellow-footed antechinus | Tasmanian devil | Gray short-tailed opossum | Agile gracile mouse opossum | European shrew | White-toothed pygmy shrew | Horse | Plains zebra | Human | Chimpanzee | House mouse | Norway rat |

Ratio 25 50 75

Figure 8 shows the graph of the 10 most enriched “biological processes” among the GO terms, except for “Negative regulation of transcription by RNA polymerase II” as its inclusion would have made the graph too large to read. The enriched terms are shown in yellow. Six of them describe a “developmental process”. The remaining three form a chain on the right hand side of the graph and are “multicellular organismal processes” that are part of “multicellular organism development”. The remaining term, “Negative regulation of transcription by RNA polymerase II”, fits the fact that development is largely based on transcriptional regulation. The Bonferroni-corrected significance of these enrichment ratios was determined by Monte Carlo simulation and is less than 0.003. This *P* -value is an upper bound that is contingent on the number of iterations in the Monte Carlo test. We used one million iterations, a greater number would have reduced the *P* -values. To gain some intuition for the true *P* -values, we plot the null distribution used in the Monte Carlo tests in human for the two GO terms “Anterior/Posterior pattern specification” and “Chordate embryonic development”, which are the top and bottom terms in Table 1. Figure 9A shows the null distribution of the number of genes found for “Anterior/Posterior pattern specification” compared to the observed value, 88. Clearly, the null distribution and the observed value are well separated. Similarly, Figure 9B shows the null distribution of the number of genes found for “Chordate embryonic development”, compared to the observed value, 159. This time the upper bound of the null distribution is even further removed from the observed value, implying an even smaller true *P* -value.

**Figure 8.**
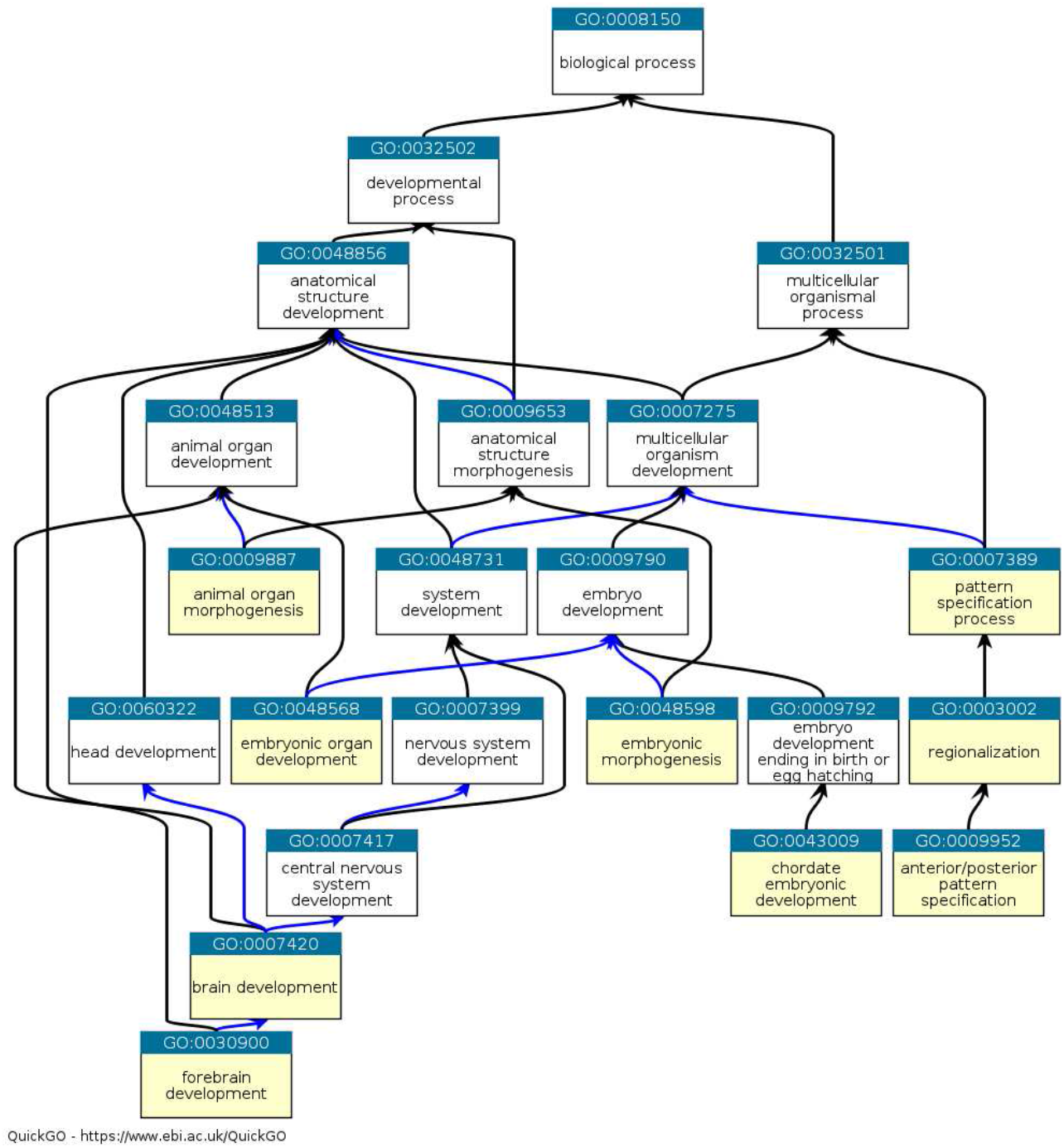
The graph of the ten most enriched GO terms listed in Table 1, except for “Negative regulation of transcription by RNA polymerase II”, for genes whose promoters intersect unique regions in mammalian genomes; enriched terms are in yellow, black arrows indicate an “is a” relationship, blue arrows a “part of” relationship.

**Figure 9.**
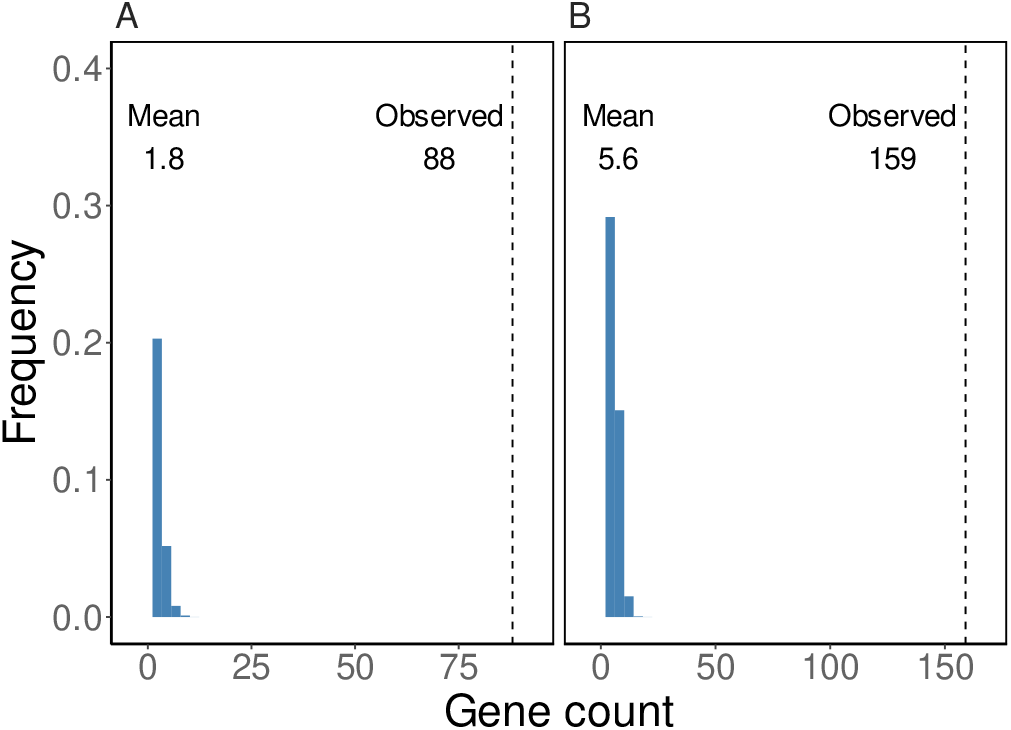
Null distribution of the number of human genes associated with the GO terms “Anterior/posterior pattern specification” (A) and “Chordate embryonic development” (B), compared to the number of genes observed indicated by the dashed vertical lines

### E. Anonymous URs

Anonymous URs are those that intersect neither known promoters nor transcripts. In our final analysis we concentrated on the longest anonymous UR per organism. These ranged from 83 kb in the tasmanian devil to 10 kb in the european shrew (Table S2). To investigate their functional content, we used Blastx to compare them to the SwissProt database. This resulted in four hits with *E <* 10^−5^ (not shown). Two of these were against LINE-1 elements, one was to the guanine nucleotide exchange factor in an unplaced contig of the genome of the gray short-tailed opossum. However, the gray short-tailed opossum has a canonical version of the gene encoding this protein on chr6:167,317,046–167,496,814. The most significant hit we found (*E* = 1.8 × 10^−13^) was between the tasmanian devil UR and human inositol polyphosphate-5-phosphatase A, INPP5A.

INPP5A is one of ten mammalian inositol 5-phosphatases, enzymes that play a crucial role in intracellular signaling (Pirruccello and De Camilli, 2012). INPP5A is special among the inositol 5-phosphatases, as it acts on soluble rather than membrane-bound inositol polyphosphates. Of the 18 mammalian genomes we studied, 17 have an annotated INPP5A. Given our Blastx results, the most likely explanation for the apparent absence of INPP5A from the genome of the tasmanian devil is that its annotation is incomplete in this respect.

## Discussion

Unique regions, URs, were of immediate interest at the beginning of the era of mammalian genomics (Simons et al., 2005). As noted at the time, URs are enriched for developmental genes and their chromatin is enriched for bivalent markings, and thus silenced but poised for activation (Bernstein et al., 2006). In the years since, RepeatMasker tracks have become a standard feature of genome browsers, and chromatin marks are widely studied as a crucial link between genotype and phenotype. However, URs have not been investigated much since then. The aim of our project was to revive the study of URs by providing simple and efficient tools for their detection and annotation, and by analyzing URs in 18 mammalian genomes (Figure 4).

As to detecting URs, we used a published tool, Macle, to calculate the match complexity, *C*_m_, which has an expectation of 1 in DNA regions that cannot be distinguished form random (Pirogov et al., 2019). Such regions have no close homologs in the rest of the genome, they are unique.

Since the properties of such regions are not as intuitive as, say, the notion of missing Blast hits, we compared the alignment-based tool RepeatMasker with Macle. Being alignment-free makes Macle much faster than RepeatMasker, but also less sensitive (Figure 2). As a result, the regions identified are only free of recent transposon insertions, while transposons with a divergence greater than 0.2 escape detection (Figure 3). Moreover, the *C*_m_ is only based on repeats, not specifically on a particular type of repeat like transposons, which in the human genome meant that a singleton transposon, HERV-Fc2, was also counted as unique, while RepeatMasker flagged it.

The classical separation between indexing and searching implemented in Macle is a standard in string-based bioinformatics tools like Blast and fast read mappers. In the case of Macle, indexing allows the sliding window analysis to run in seconds, while indexing takes roughly an hour (Figure 5A). Similarly, the memory consumption of indexing is roughly two orders of magnitude greater than that of the sliding window analysis (Figure 5B).

The sliding window analysis is highly sensitive to window length, where an increase leads to a decrease in yield (Figure 6A). This corresponds to our intuition that a long region is less likely to escape transposon insertions over long periods of time than a short region. With decreasing yield the proportion of intersecting promoters also decrease from more than half to zero (Figure 6B). Similarly, the proportion of enriched biological processes declines from over 30 % to zero with increasing window length (Figure 6C).

For each of the nine mammalian orders we investigated, the two genomes sampled closely tracked each other’s graph of the average enrichment ratio as a function of window length (Figure 7). Given that the longest URs contain loci like the *Hox* clusters, that is, a large number of genes of one narrow functional category, we expected a steady increase in enrichment ratio as a function of window length. However, across the orders the trajectories varied between plateauing in most orders, and steadily increasing in primates and shrews (Eulipotyphla).

Still, the classical observation that URs are enriched for developmental genes is true for all 18 taxa investigated (Table 1, Figure 8). Now, URs are not the only sequence property associated with function, CpG islands are a classical sequence feature associated with transcription initiation in vertebrates (Deaton and Bird, 2011). It was previously found that in human and mouse 88 % of URs at least 10 kb long intersect a CpG island (Pirogov et al., 2019). This large overlap between the two features might suggest that URs are essentially CpG islands. However, the specific enrichment for developmental genes observed in URs is only true for CpG islands longer than 2 kb (Elango and Yi, 2011). The overlap between URs and long CpG islands is much smaller than between URs and CpG islands in general. In human, 35 % of URs intersect CpG islands of 2 kb or more, in mouse this fraction is down to 19 % (Pirogov et al., 2019).

We called URs without any intersecting promoters or transcripts “anonymous”. As a quick check of their functional content, we compared them to SwissProt, a carefully curated sample of proteins. Our expectation was that we would find nothing, annotation by homology being a standard component of annotation pipelines. Nevertheless, in the 83 kb UR of the tasmanian devil we found a hit to the signaling enzyme inositol polyphosphate-5-phosphatase A (INPP5A). This enzyme is known in the other 17 genomes investigated. We are thus confident that we have located an essential, albeit hitherto missing, gene in the genome of the tasmanian devil.

Our analysis of anonymous URs only scratches the surface of what could be done, including annotation by nucleotide Blast instead of protein Blast, and the identification of transcription factor binding sites (Bailey et al., 2015).

We conclude that Macle makes it easy to find URs on various scales and that the workflow implemented in the Augre repository allows their efficient annotation and enrichment analysis. The importance of long URs is underscored by the general observation that they are highly enriched for developmental genes, and on the specific discovery that the longest anonymous UR in the tasmanian devil apparently contains a component of intracellular signaling.

## Author contributions statement

B.V.M. and B.H. designed the study. B.V.M. carried out the data analysis. B.V.M. and B.H. wrote the manuscript.

## Data availability

All relevant data for this study is available from the dataverse at doi.org/10.17617/3.4IKQAG.

