## Supplementary material for "Detection and Annotation of Unique Regions in Mammalian Genomes": Tables S1 and S2

Table S1: Genome accessions, corresponding mammalian order for each species, total genome size (Gb).

| # | Accession | Order | Species | Common name | Size (Gb) |
|---|---|---|---|---|---|
| 1 | GCF_002263795.2 | Artiodactyla | *Bos taurus* | Bovine | 2.7 |
| 2 | GCF_000003025.6 | Artiodactyla | *Sus scrofa* | Pig | 2.5 |
| 3 | GCF_011100685.1 | Carnivora | *Canis lupus familiaris* | Domestic dog | 2.5 |
| 4 | GCF_016509475.1 | Carnivora | *Prionailurus bengalensis* | Leopard cat | 2.4 |
| 5 | GCF_004126475.2 | Chiroptera | *Phyllostomus discolor* | Pale spear-nosed bat | 2.1 |
| 6 | GCF_022682495.1 | Chiroptera | *Desmodus rotundus* | Common vampire bat | 2.1 |
| 7 | GCF_016432865.1 | Dasyuromorphia | *Antechinus flavipes* | Yellow-footed antechinus | 3.2 |
| 8 | GCF_902635505.1 | Dasyuromorphia | *Sarcophilus harrisii* | Tasmanian devil | 3.1 |
| 9 | GCF_000002295.2 | Didelphimorphia | *Monodelphis domestica* | Gray short-tailed opossum | 3.6 |
| 10 | GCF_016433145.1 | Didelphimorphia | *Gracilinanus agilis* | Agile gracile mouse opossum | 3.7 |
| 11 | GCF_024139225.1 | Eulipotyphla | *Suncus etruscus* | White-toothed pygmy shrew | 2.5 |
| 12 | GCF_027595985.1 | Eulipotyphla | *Sorex araneus* | European shrew | 2.4 |
| 13 | GCF_002863925.1 | Perissodactyla | *Equus caballus* | Horse | 2.5 |
| 14 | GCF_021613505.1 | Perissodactyla | *Equus quagga* | Plains zebra | 2.5 |
| 15 | GCF_000001405.40 | Primates | *Homo sapiens* | Human | 3.3 |
| 16 | GCF_028858775.1 | Primates | *Pan troglodytes* | Chimpanzee | 3.2 |
| 17 | GCF_000001635.27 | Rodentia | *Mus musculus* | House mouse | 2.7 |
| 18 | GCF_015227675.2 | Rodentia | *Rattus norvegicus* | Norwegian rat | 2.6 |

Table S2: The longest anonymous regions in each organism investigated; chromosome *u* is *unplaced*.

| Assembly | Name | Chromosome | Start | End | Length |
|---|---|---|---:|---:|---:|
| GCF_902635505.1 | Tasmanian devil | NC_045427.1 (2) | 660,845,372 | 660,928,371 | 83,000 |
| GCF_016433145.1 | Agile gracile mouse opossum | NC_058136.1 (X) | 61,932,012 | 61,993,011 | 61,000 |
| GCF_022682495.1 | Common vampire bat | NC_071393.1 (7) | 1,735,042 | 1,795,041 | 60,000 |
| GCF_016432865.1 | Yellow-footed antechinus | NC_067404.1 (X) | 72,884,124 | 72,938,123 | 54,000 |
| GCF_004126475.2 | Pale spear-nosed bat | NC_040917.2 (15) | 25,189,025 | 25,238,024 | 49,000 |
| GCF_016509475.1 | Leopard cat | NC_057358.1 (B4) | 140,638,285 | 140,675,284 | 37,000 |
| GCF_000002295.2 | Gray short-tailed opossum | NW_001583633.1 (u) | 31,561 | 65,560 | 34,000 |
| GCF_011100685.1 | Domestic dog | NC_049238.1 (17) | 1,452,570 | 1,482,569 | 30,000 |
| GCF_000003025.6 | Pig | NC_010453.5 (11) | 77,807,181 | 77,837,180 | 30,000 |
| GCF_028858775.1 | Chimpanzee | NC_072412.1 (14) | 92,813,178 | 92,840,177 | 27,000 |
| GCF_021613505.1 | Plains zebra | NC_060274.1 (8) | 129,719,911 | 129,745,910 | 26,000 |
| GCF_002263795.2 | Bovine | NW_020192290.1 (u) | 169,985 | 193,984 | 24,000 |
| GCF_024139225.1 | White-toothed pygmy shrew | NC_064855.1 (8) | 123,975,861 | 123,996,860 | 21,000 |
| GCF_000001405.40 | Human | NC_000004.12 (4) | 10,214,304 | 10,235,303 | 21,000 |
| GCF_002863925.1 | Horse | NC_009175.3 (X) | 22,668,551 | 22,686,550 | 18,000 |
| GCF_000001635.27 | House mouse | NC_000086.8 (X) | 56,958,927 | 56,971,926 | 13,000 |
| GCF_015227675.2 | Norwegian rat | NC_051339.1 (4) | 123,771,487 | 123,783,486 | 12,000 |
| GCF_027595985.1 | European shrew | NC_073302.1 (1) | 164,574,001 | 164,584,000 | 10,000 |
